# Effect of biomaterial stiffness on cardiac mechanics in a biventricular infarcted rat heart model with microstructural representation of *in situ* intramyocardial injectate

**DOI:** 10.1101/2022.05.23.493036

**Authors:** YD Motchon, KL Sack, MS Sirry, M Kruger, E Pauwels, D Van Loo, A De Muynck, L Van Hoorebeke, NH Davies, T Franz

## Abstract

Intramyocardial delivery of biomaterials is a promising concept for treating myocardial infarction. The delivered biomaterial provides mechanical support and attenuates wall thinning and elevated wall stress in the infarct region. This study aimed at developing a biventricular finite element model of an infarcted rat heart with a microstructural representation of an *in situ* biomaterial injectate, and a parametric investigation of the effect of the injectate stiffness on the cardiac mechanics.

A three-dimensional subject-specific biventricular finite element model of a rat heart with left ventricular infarct and microstructurally dispersed biomaterial delivered one week after infarct induction was developed from *ex vivo* microcomputed tomography data. The volumetric mesh density varied between 303 mm^-3^ in the myocardium and 3,852 mm^-3^ in the injectate region due to the microstructural intramyocardial dispersion. Parametric simulations were conducted with the injectate’s elastic modulus varying from 4.1 to 405,900 kPa, and myocardial and injectate strains were recorded.

With increasing injectate stiffness, the end-diastolic median myocardial fibre and cross-fibre strain decreased in magnitude from 3.6% to 1.1% and from −6.0% to −2.9%, respectively. At end-systole, the myocardial fibre and cross-fibre strain decreased in magnitude from −20.4% to −11.8% and from 6.5% to 4.6%, respectively. In the injectate, the maximum and minimum principal strains decreased in magnitude from 5.4% to 0.001% and from −5.4% to −0.001%, respectively, at end-diastole and from 38.5% to 0.06% and from −39.0% to −0.06%, respectively, at end-systole.

With the microstructural injectate geometry, the developed subject-specific cardiac finite element model offers potential for extension to cellular injectates and *in silico* studies of mechanotransduction and therapeutic signalling in the infarcted heart with an infarct animal model extensively used in preclinical research.

## 1. Introduction

Cardiovascular disease (CVD) is the leading cause of death worldwide. In 2017, CVD was responsible for approximately 17.8 million deaths (Kaptoge *et al*. 2019, Mendis *et al*. 2011). An increase in CVD-related deaths by 26.9%, from 17.5 million (i.e. 31% of global deaths) in 2012 to 22.2 million by 2030, has been predicted. Alarmingly, in the working-age population of low- and middle-income countries, including South Africa, the rate of people affected is becoming considerably high (Finegold *et al*. 2013, Kaptoge *et al*. 2019, Opie and Mayosi 2005). This situation affects the global economy and therefore influences social cohesion in the communities.

Myocardial infarction (MI) originates from coronary occlusion, causing a lack of oxygenated blood supply to a specific myocardial region. In the long term, cardiomyocyte death is followed by scar formation, causing excessive load to heart dysfunction and heart failure. Current treatments for MI include reperfusion (drug administration or surgery), mechanical device assistance (e.g. left ventricle assist device) and heart transplantation. New therapy approaches based on biomaterial injections have emerged. Clinical studies involving such therapeutic biomaterials showed promising results addressing adverse ventricular remodelling following post-MI inflammatory response (Christman *et al*. 2004, Fan *et al*. 2019, *Johnson and Christman 2013, Silveira-Filho *et al*. 2021)*.

Limitations of experimental studies are the cost of resources and the invasiveness of *in vivo* protocols. Computational modelling appeared as an alternative to investigate the mechanical mechanisms and parameters involved in therapeutic biomaterial injections in infarcted hearts, such as biomaterial stiffness, delivery location, injection volume, and injection pattern (Cai *et al*. 2017, Kortsmit *et al*. 2013a, Kortsmit *et al*. 2013b, Wall *et al*. 2006, Wang *et al*. 2017, Wenk *et al*. 2011, Wenk *et al*. 2009, Wise *et al*. 2016).

Wall *et al*. (2006) reported a reduction in the wall stress, affecting the ventricular function, using an ovine heart finite element (FE) model with a non-contractile biomaterial injectate in the left ventricular (LV) wall implemented by modification of the FE mesh. The results depended on the volume, the location, and the material stiffness of the injected biomaterial.

The morphology and dispersion of the intramyocardial injectates are challenging to control. Wang *et al*. (2017) studied the impact of injectate volume and stiffness on LV myofiber stress and wall thickness. They showed that a larger volume and stiffer material contributes to myofibre stress reduction and wall thickness increase in the LV, which is essential to alter ventricular remodelling. Computational models have been used to optimize the pattern (Sepantafar *et al*. 2016, Wenk *et al*. 2009) and the volume of biomaterial injectates (Wise *et al*. 2016).

The current study aimed to develop a biventricular finite element model of a rat heart with an antero-apical infarct and an intramyocardial biomaterial injectate delivered in the infarct region one week after infarct induction. A parametric study was undertaken to investigate the effect of injectate stiffness on cardiac mechanics. Emphasis was placed on the realistic microstructural geometrical representation of the *in situ* intramyocardial dispersion of the biomaterial injectate. The intramyocardial injectate constitutes an important configuration to investigate the mechanical and mechanotransductory responses of therapeutic cells transplanted with the biomaterial for the treatment of myocardial infarction.

## 2. Materials and methods

### 2.1 Volumetric image data of infarcted rat heart with intramyocardial injectate

*Ex vivo* microcomputed tomography (µCT) image data of an infarcted rat heart with polymeric intramyocardial injectate from an unrelated study (unpublished data) were used for geometric reconstruction. In brief, male Wistar rats (body mass: 180-220 g) were anaesthetized, and the heart was exposed via left thoracotomy along the 4^th^ intercostal space. Myocardial infarction was induced by permanent ligation of the left anterior descending coronary artery 3 mm distal to the auricular appendix. The discolouration of the anterior ventricular wall and reduced contractility were hallmarks of a successful occlusion of the artery. The chest was stepwise closed, and buprenorphine was administered for pain management. Seven days later, the heart was accessed via the 4^th^ intercostal space and 100 µl of radiopaque silicone rubber containing lead chromate (Microfil® MV-120 Flow-Tech, Carver, MA, USA) diluted 1.5:1 with MV-diluent was injected into the infarct area. Dispersion and *in situ* polymerization of the Microfil® material were allowed for 30 min after the injection. The animals were then humanely killed, and the hearts were carefully harvested, thoroughly rinsed with saline, fixed in a 4% paraformaldehyde solution and transferred to saline for µCT scanning. All animal experiments were authorized by the Institutional Review Board of the University of Cape Town and performed according to the National Institutes of Health (NIH, Bethesda, MD, USA) guidelines.

The µCT scans were performed with a custom-made scanner with a Feinfocus X-ray tube and a Varian 2520V Paxscan a-Si flat panel detector (CsI screen, 1920 × 1536, 127 µm pixel size) at the Centre for X-ray Tomography of Ghent University (UGCT) (Masschaele *et al*. 2007). For each scan, 1,801 projections were captured with an exposure time of 0.8 s. The resulting scan images had a voxel pitch of 10 µm. Reconstruction was performed using the UGCT software package Octopus (Vlassenbroeck *et al*. 2007).

### 2.2 Three-dimensional reconstruction and meshing of a biventricular cardiac geometry

Only the two ventricles were considered for the FE model, resulting in a biventricular (BV) geometry truncated at the base. Before segmentation, the orientation of the image stack was aligned with the longitudinal cardiac axis. The image segmentation involved region-growing, level-set thresholding and manual segmentation (Simpleware ScanIP, Synopsys). Two masks were created, distinguishing the cardiac tissue and the injectate.

The resulting geometry captured the essential morphology of the left (LV) and right ventricle (RV) and the dispersed intramyocardial injectate in the LV free wall (Figure 1 a and Table 1). The geometry was meshed with 206,142 quadratic tetrahedral elements (injectate: 58,902 elements, myocardium: 147,240 elements). The mesh density varied between 302.8 mm^-3^ in the myocardium and 3,852.3 mm^-3^ in the injectate region (Figure 1 c).

**Figure 1.**
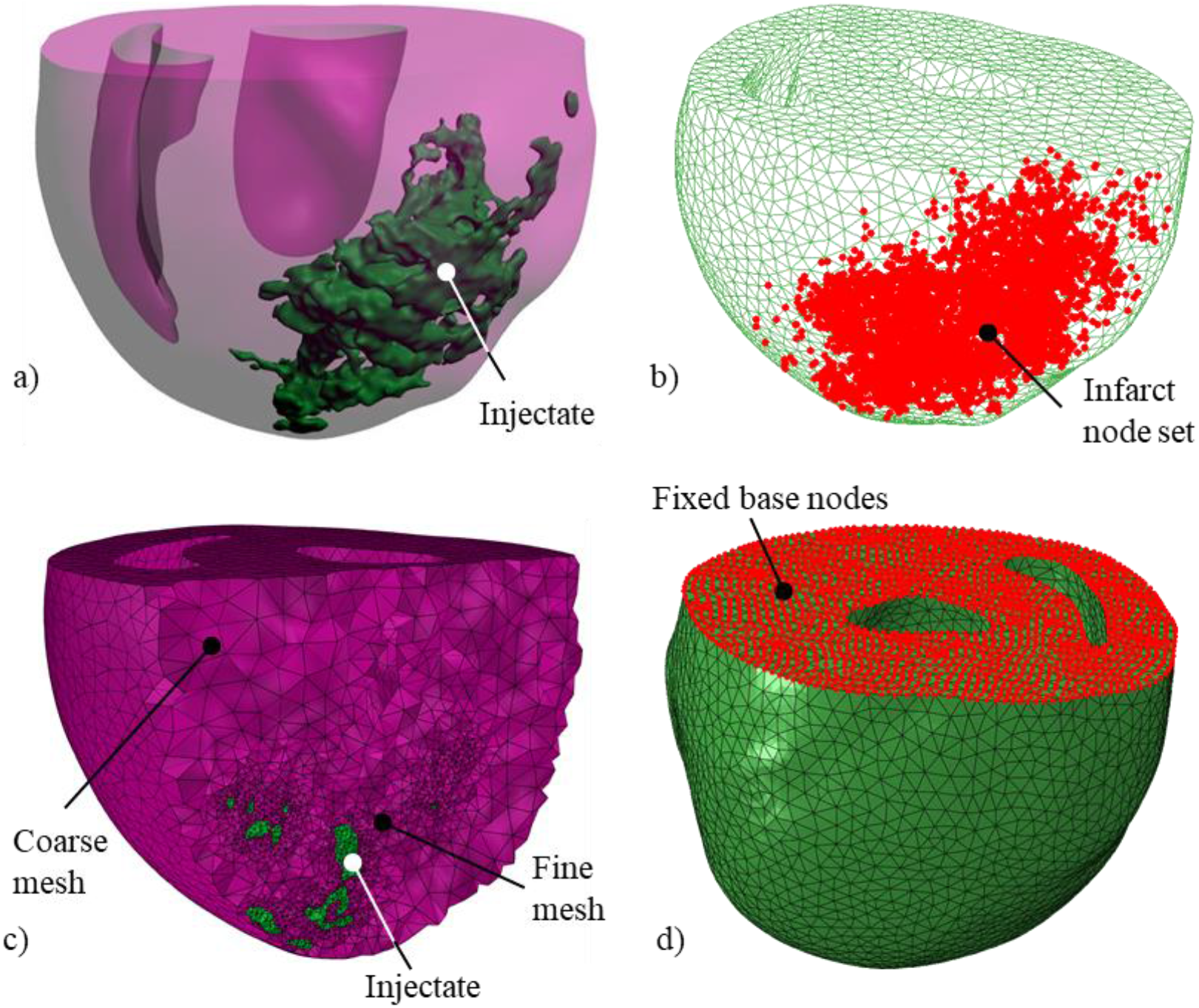
a) Biventricular cardiac geometry of the rat heart developed from μCT image data with LV, RV, and dispersed intramyocardial injectate in the infarcted region of the LV. b) Infarct region shown with nodes set, the infarct was defined around the injectate. c) Increase in mesh density from the myocardium to the injectate to accommodate the microstructural dispersion of the injected material. d) Nodes on base with boundary condition of zero displacement in the longitudinal direction to prevent rigid body motion (red points).

**Table 1.**
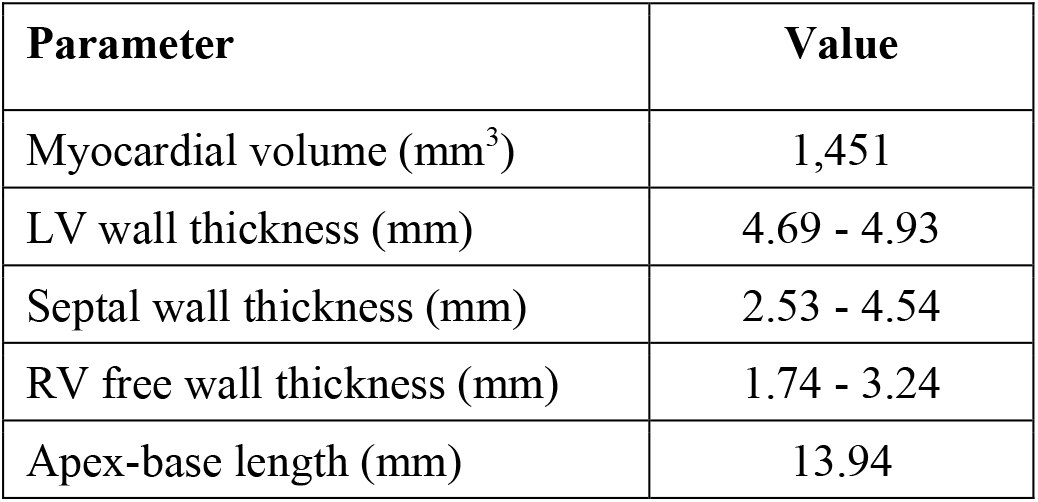
Morphometric data of the biventricular geometry of the rat heart.

| Parameter | Value |
| --- | --- |
| Myocardial volume ($\text{mm}^3$) | 1,451 |
| LV wall thickness (mm) | 4.69 - 4.93 |
| Septal wall thickness (mm) | 2.53 - 4.54 |
| RV free wall thickness (mm) | 1.74 - 3.24 |
| Apex-base length (mm) | 13.94 |

The meshed geometry was imported in Abaqus 6.14-3 CAE (Dassault Systèmes, Providence, RI, USA). After that, the infarct was approximated in the antero-apical region of the LV and defined as a node set.

### 2.3 Finite element model development

The meshed BV geometry was imported in Abaqus CAE (Abaqus 6.14-3, Dassault Systèmes, Providence, RI, USA) to implement the different mechanical properties and investigate cardiac tissue mechanics. A 10-node tetrahedral element type (C3D10M) was used to obtain a discretized FE problem.

#### 2.3.1 Myofibre structure

A rule-based approach was used to describe the myofibre angle distribution in the myocardium. A Matlab code developed by Sack *et al*. (2018) was used to generate fibre orientation values through the myocardium. The primary function of the algorithm was to find the different components of the fibre vector within each element in the meshed region of the myocardium. The projected values of the fibre vector were obtained with two principal angles. The helix angle α_h_, is formed by the projection of the fibre on the plane created by the circumferential-longitudinal unit vector (u_c_, u_l_) and the circumferential unit vector u_c_. The transverse angle α_t_ between the projection of the fibre on the plane is formed by the radial-circumferential unit vector (f_p1_) and the circumferential unit vector. In the current study, the fibre angle was −50° to 80° from the epicardial to the endocardial surfaces (Chen *et al*. 2003, Sirry 2015). The obtained fibre orientation data were incorporated in the FE model in Abaqus.

#### 2.3.2 Constitutive laws

##### Healthy and infarcted myocardium

The passive mechanical properties of the myocardium were described with a hyperelastic anisotropic law using a modified strain energy function from Holzapfel and Ogden (2009). The changes were introduced to consider the pathological stage of the heart tissue (Sack *et al*. 2018). The passive mechanical properties of the infarcted myocardium depend on the stage of the infarct (Holmes *et al*. 2005, Holmes *et al*. 1997, Tyberg *et al*. 1974, Villarreal *et al*. 1991). A one-week infarct stage was considered in the current study. Hence, an increase in stiffness (Hood *et al*. 1970) in the fibre, circumferential and longitudinal direction was implemented with the parameters h and p in the Eqn. (2), with h = 1 and 0 representing a healthy and infarcted myocardium, respectively.

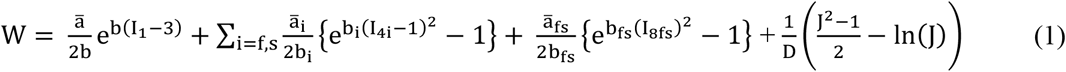

with

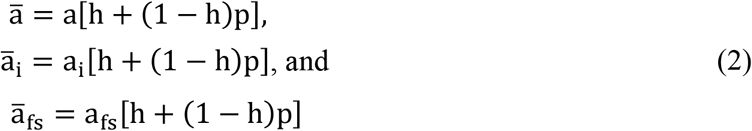

The active contraction of the myocardium was implemented with a time-varying elastance approach (Guccione and Mcculloch 1993, Guccione *et al*. 1993, Walker *et al*. 2005), with addition of tissue health parameter h to represent the pathological stage of the myocardium (Sack *et al*. 2018):

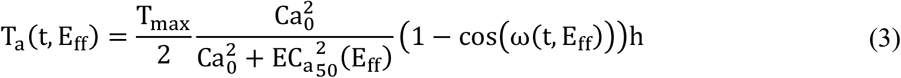

with

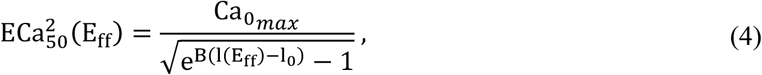

and

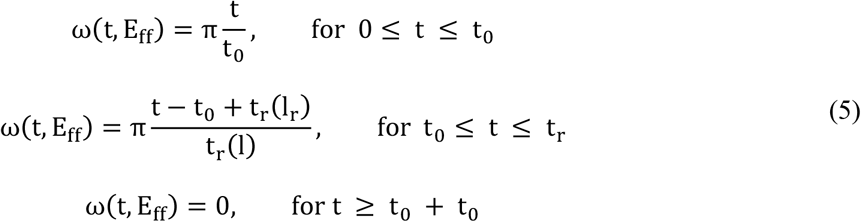

The additive approach was used to determine the total tension in the myocardium where, the time-varying active tension T_a_ (Eqn (3)) was combined with the tension derived from the passive response, T_p_:

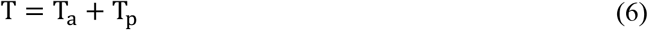

The description and values of the passive and active constitutive parameters are provided in Supplemental Tables 1 and 2.

##### Injectable biomaterial

The injectable biomaterial, e.g. polyethylene glycol (PEG) hydrogel, was described as hyperelastic isotropic incompressible material with a Neo-Hookean material model:

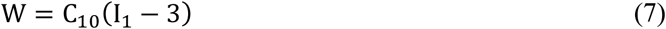

where I_1_ is the first deviatoric strain invariant, and C_10_ characterizes the material stiffness obtained from the elastic modulus E_inj_ and the Poisson’s ratio ν_inj_:

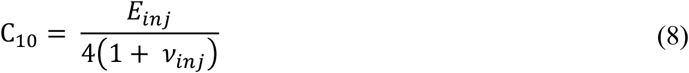

with ν_inj_ = 0.5 to represent incompressibility and E_inj_ between 4.1 and 405,900 kPa (see section 2.4 Finite element simulations for further details).

#### 2.3.3 Boundary conditions

A zero displacement was applied to the nodes at the base in the longitudinal direction to prevent the rigid body motion of the geometry, see Figure 1(d). The passive filing was implemented by a linearly increasing pressure load on both LV and RV cavity surfaces. Several studies reported a higher cavity pressure in the LV than in the RV. In normal human hearts, systolic pressure was 30-40 mmHg in the RV and 100-140 mmHg in the LV (Mininni *et al*. 1996, Reichek and Devereux 1982). Pacher *et al*. (2004) measured left ventricular end-diastolic and end-systolic pressure in rats and found 3.8 ± 0.9 mmHg and 133.8 ± 8.1 mmHg, respectively. In the current study, the cavity pressure was taken from 0 to 3.0 mmHg for the LV and from 0 to 0.75 mmHg for the RV. This choice agrees with the range of the experimental end-diastolic pressure findings.

#### 2.3.4 Determination of computation time for end-systole and end-diastole

The end-diastolic time point was determined by applying a linearly increasing pressure on the LV and RV endocardial surfaces until the LV cavity volume matched a target experimental ED volume by Pacher *et al*. (2004). This step was performed on the BV model with injectate elastic modulus E_inj_ = 73.8 kPa.

A second simulation was performed to determine the end-systolic time point. The obtained time corresponded to the contraction and was determined from the ED time point defined previously until the active tension declined. The LV volume was calculated at this time point and compared with end-systolic volume values in the literature (Pacher *et al*. 2004).

### 2.4 Finite element simulations and data acquisition

A parametric study to determine the impact of the injectate stiffness on the deformation of the myocardium and the biomaterial injectate was conducted with a range of values for the elastic modulus of injectate, i.e. E_inj_ = 4.1, 7.4, 40.6, 73.8, 405.9, 738, 4,059, 7,380, 40,590, 73,800 and 405,900 kPa.

The strain in the myocardium and the biomaterial injectate was recorded at the end-diastolic and end-systolic time points of the cardiac cycle for each E_inj_ value. The myofibre and cross-fibre strains were used for the healthy and infarcted myocardium, whereas the maximum principal and minimum principal strains were used for the injectate. The *ex vivo* geometry of the heart developed from μCT image data was used as reference configuration for strain calculations. This configuration was assumed unloaded, and pre-stress was not considered.

For each finite element in the mesh, the element strain, ε_EL_, was determined as the arithmetic mean of the strain values at the four integration points, ε_IP,i_, of the 10-node tetrahedral elements.

### 2.5 Statistical analysis

Descriptive statistical analysis was performed on the strain data to determine the normality (Shapiro-Wilk normality test) and variability (SciPy, https://scipy.org/ and NumPy, https://numpy.org/). Data were presented using box and whisker plots (MatPlotLib, Python, https://matplotlib.org/) indicating median and interquartile ranges.

## 3. Results

Representative end-diastolic and end-systolic strain distributions are shown for the myofibre and cross-fibre strain in the BV wall in Figure 2 and the maximum and minimum principal strain in the injectate in Figure 3, for an elastic modulus of the injectate of E_inj_ = 73.8 kPa.

**Figure 2.**
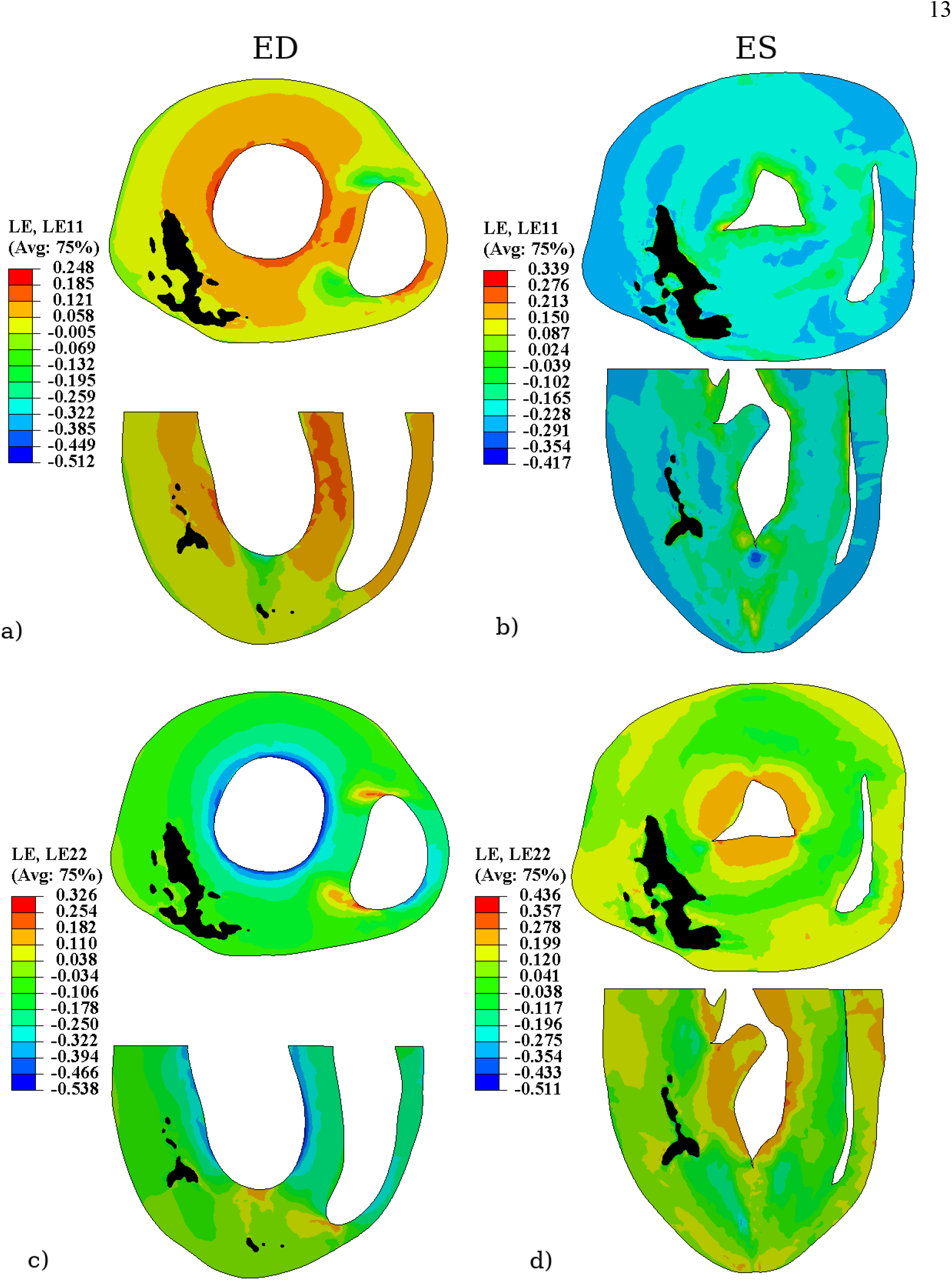
Short-axis and longitudinal contour plots showing myofibre strain (a, b) and cross-fibre strain (c, d) in the BV model at end-diastolic (ED, left column) and end-systolic time point (ES, right column) for an injectate elastic modulus E_inj_ = 73.8 kPa. The strain distribution in the injectate (black areas) is shown in Figure 3.

**Figure 3.**
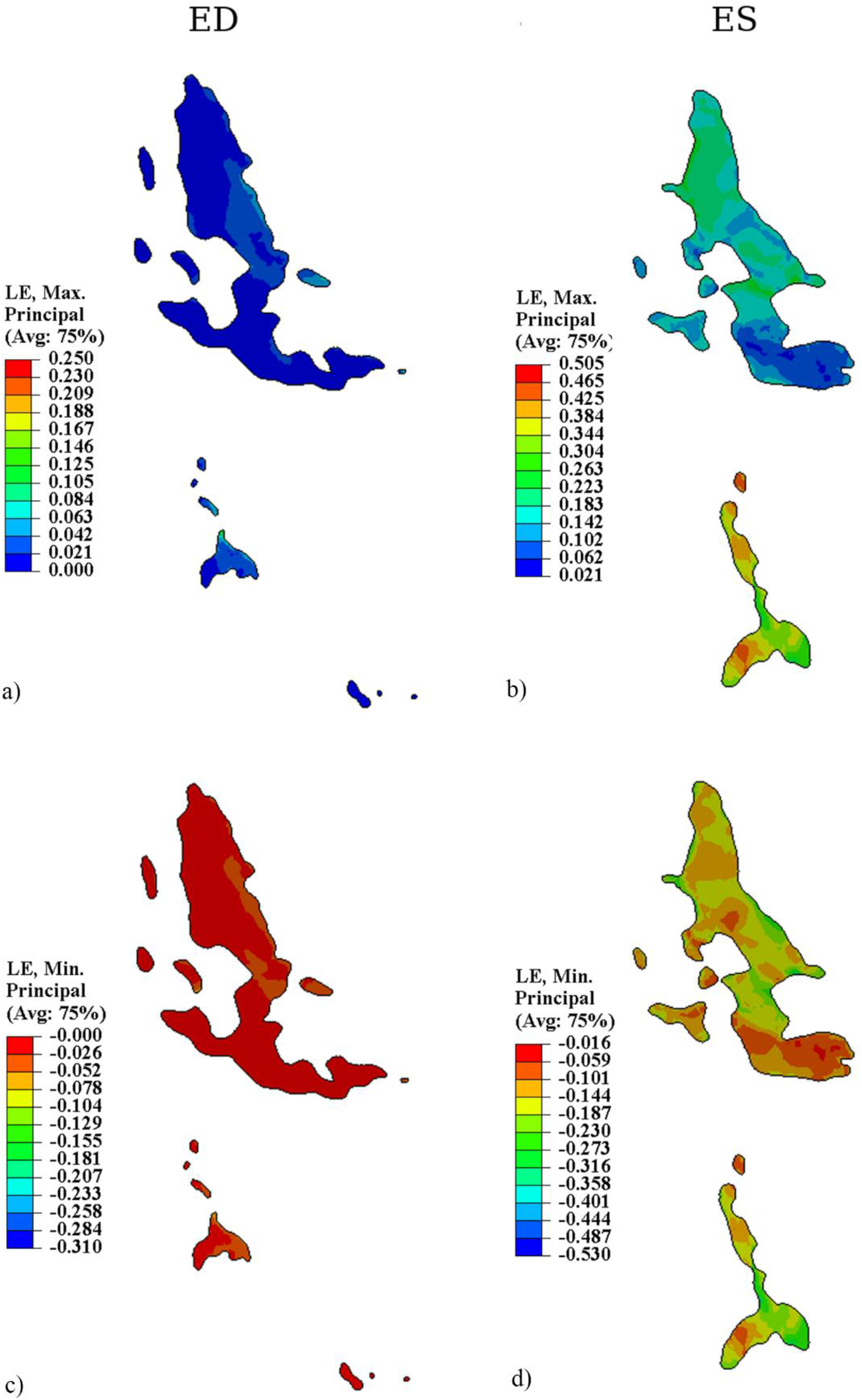
Short-axis and longitudinal contour plots of maximum (a, b) and minimum principal strain (c, d) in the injectate at end-diastolic (ED, left column) and end-systolic time point (ES, right column) for an injectate elastic modulus E_inj_ = 73.8 kPa.

### 3.1 Effect of injectate stiffness on myocardial deformation

The median end-diastolic myofibre and cross-fibre strain decreased from 3.5% to 1.0% and −5.9% to - 2.7% (decrease in magnitude) for the increase in the injectate elastic modulus (Figure 4 a and b). These changes in strain appear to be more pronounced in the low modulus region E_inj_ = 4.1 to 738 kPa and only marginal for E_inj_ > 738 kPa.

**Figure 4.**
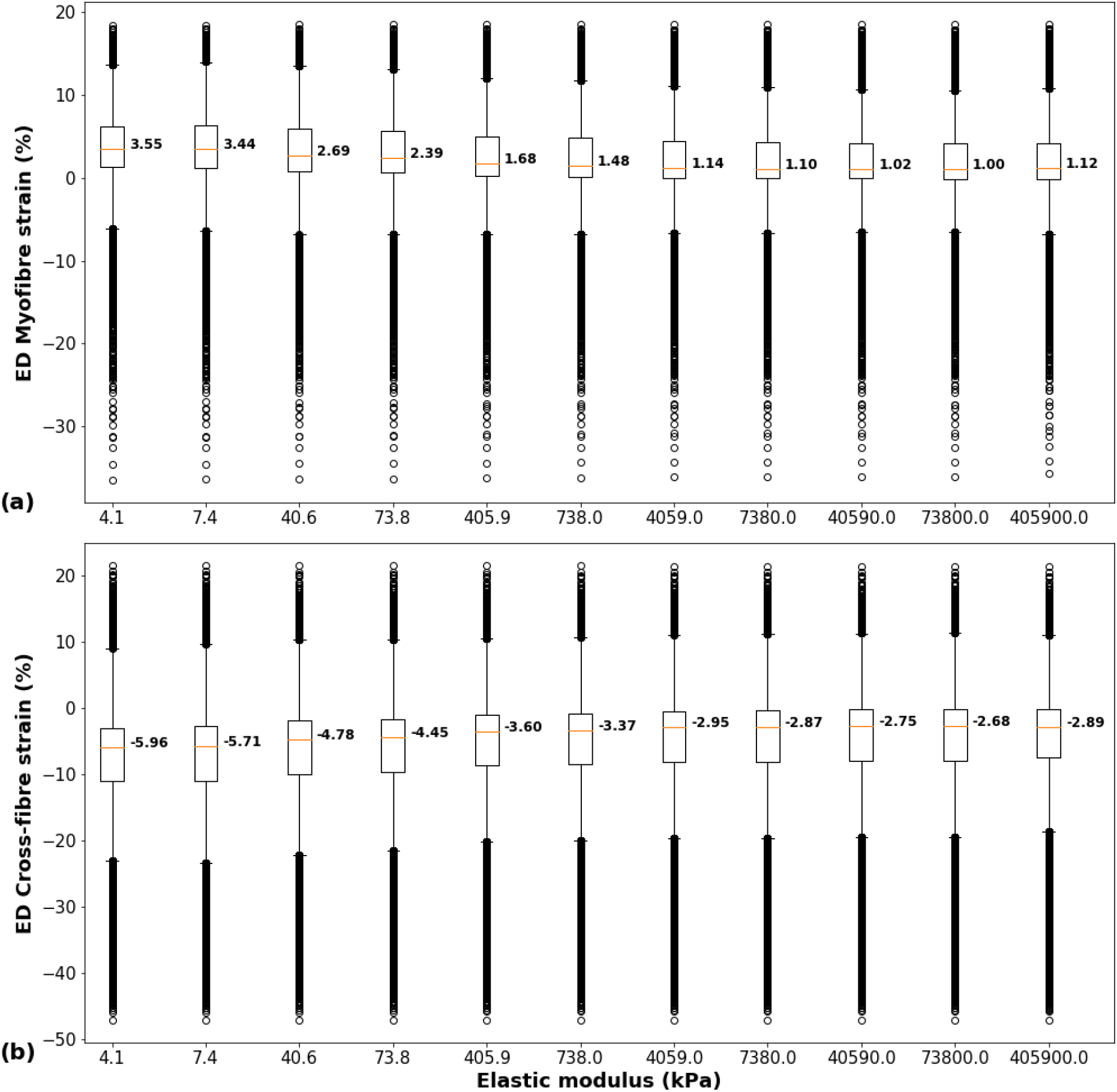
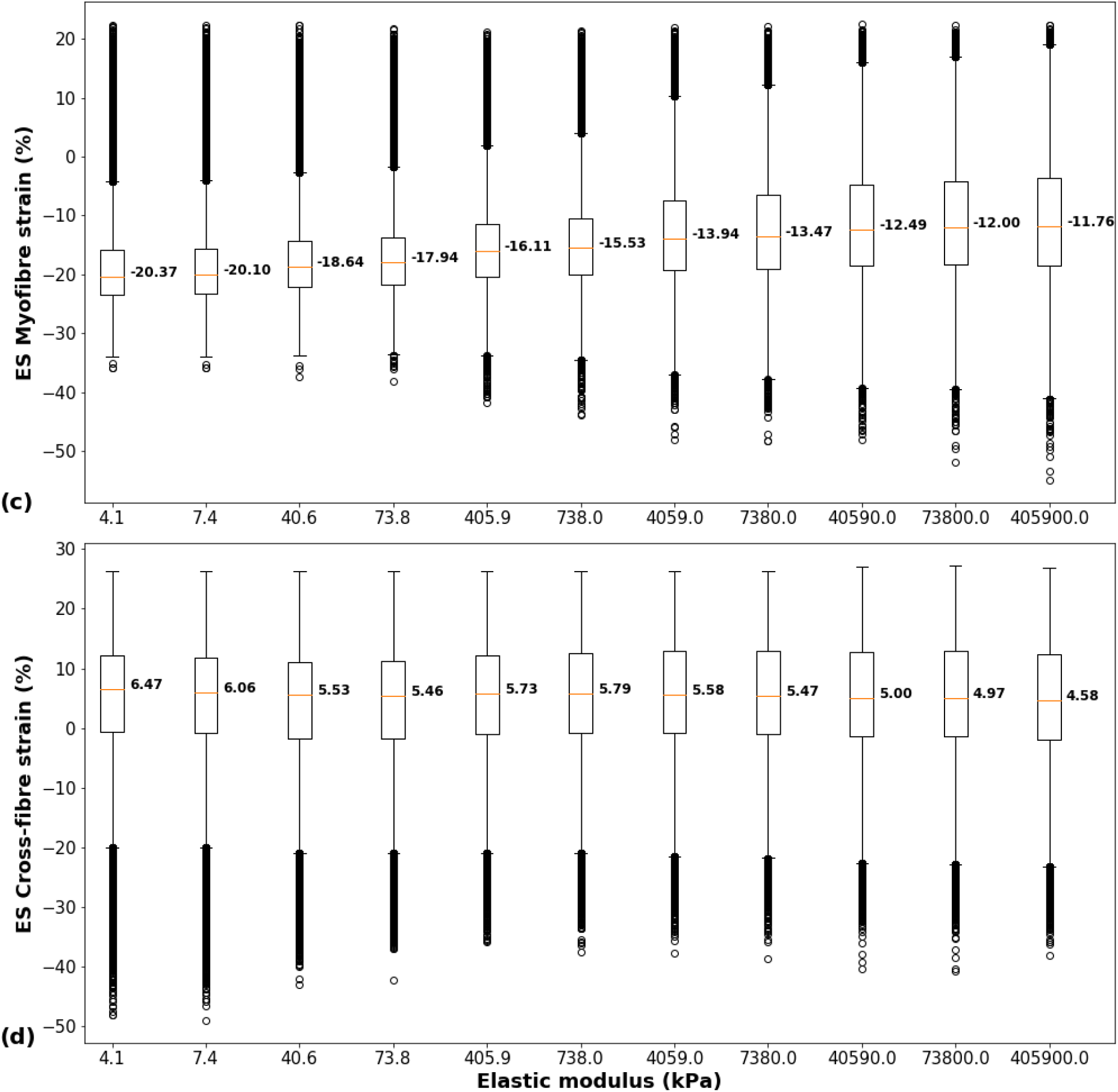
Myocardial deformation. End-diastolic (ED) myofibre (a) and cross-fibre (b) strain and end-systolic (ES) myofibre (c) and cross-fibre (d) strain versus injectate elastic modulus. The box and whiskers indicate the median (red line in box), interquartile range (IQR) from first and third quartile (lower and upper bound of the box), 1.5x IQR (lower and upper whisker), and data larger or smaller than 1.5x IQR (open circles). Each data point represents the strain value ε_EL_ in an element of the finite element mesh. Data larger or smaller than 1.5x IQR are not considered outliers but true data.

At end-systole, the median myofibre strain decreased in magnitude from −20.4% to −11.8%, with increasing elastic modulus. The median cross-fibre strain decreased from 6.5% to 4.6% with an increasing elastic modulus with an intermittent marginal increase for E_inj_ = 405.9 kPa and 738.0 kPa (Figure 4 c and d).

### 3.2 Effect of injectate stiffness on injectate deformation

As a general observation, the magnitude (i.e. median), range (difference between the highest and lowest element strain) and interquartile range of maximum and minimum principal strain decreased considerably with increasing injectate stiffness for lower E_inj_ = 4.1 - 738 kPa. However, the strain decreased marginally for the higher E_inj_ = 4,059 to 405,900 kPa, both at end-diastolic and end-systolic time points.

At end-diastole, the median maximum principal strain decreased from 5.4% to 0.001%, and the median minimum principal strain decreased in magnitude from −5.4% to −0.001% for increasing injectate stiffness (Figure 5 a and b).

**Figure 5.**
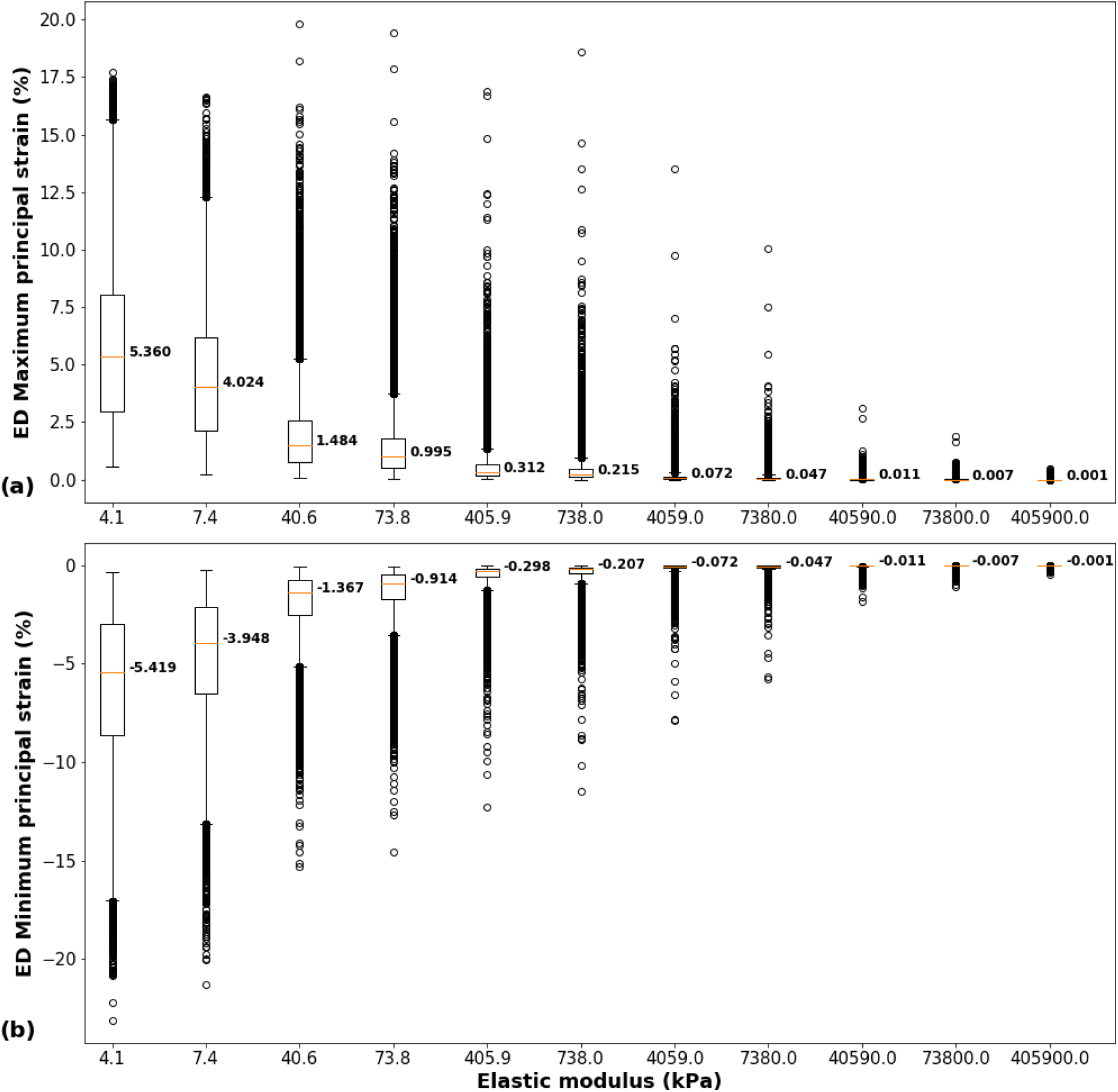
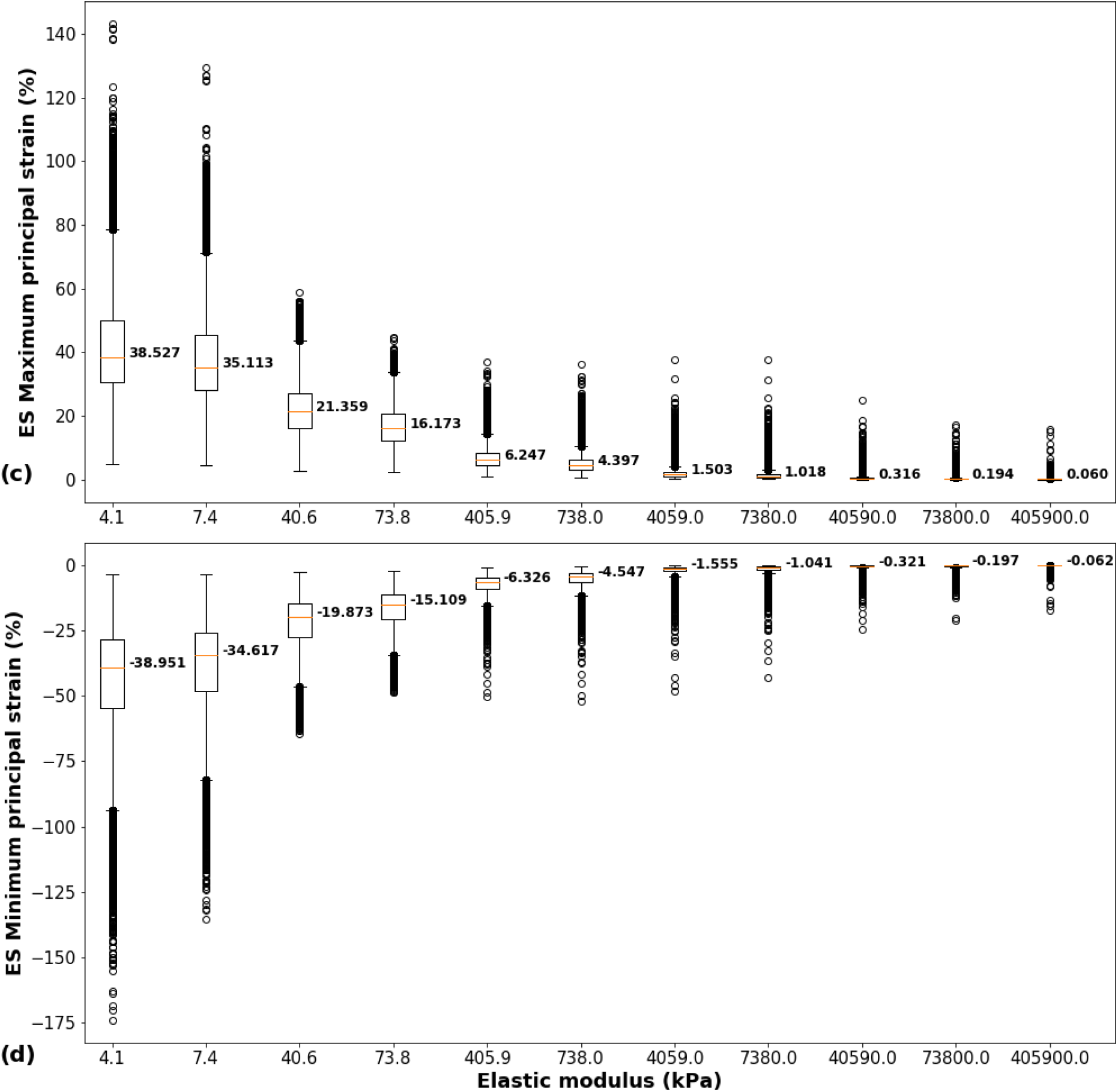
Injectate deformation. End-diastolic (ED) maximum (a) and minimum (b) principal strain and end-systolic (ES) maximum (c) and minimum (d) principal strain. The box and whiskers indicate the median (red line in box), interquartile range (IQR) from the first and third quartile (lower and upper bound of the box), 1.5x IQR (lower and upper whisker), and data larger or smaller than 1.5x IQR (open circles). Each data point represents the strain value ε_EL_ in an element of the finite element mesh. Data larger or smaller than 1.5x IQR are not considered outliers but true data.

At end-systole, the median maximum principal strain decreased from 38.5% to 0.06%, and the median minimum principal strain decreased in magnitude from −39.0% to −0.06% with increasing injectate stiffness (Figure 5 c and d).

## 4. Discussion

The present computational study involved the development of computational models for cardiac mechanics. The effect of the therapeutic biomaterial injectate stiffness on the cardiac mechanics during a cardiac cycle was shown using the developed models. At a given injectate elastic modulus, the end-diastolic myofibre strain decreased from the endocardial to the epicardial surface, which has also been reported in other studies (Costa *et al*. 1999, Takayama *et al*. 2002, Waldman *et al*. 1988). The biomaterial delivery 7 days after infarct induction was chosen since significant improvements in cardiac function and scar tissue mechanics were reported *in vivo* for the injection of PEG hydrogel one week following the infarction compared to biomaterial delivery immediately after the infarction (Kadner *et al*. 2012).

Many computational studies involving animal species used simplified cardiac geometries (Dieudonné 1969, Wall *et al*. 2006, Wenk *et al*. 2009). The current study was the first to develop a subject-specific biventricular geometry of a rat heart with *in situ* microstructural details of an intramyocardial injectate in the LV free wall. This detailed representation of the injectate is important for investigating the mechanics and mechanotransduction of cells delivered with and embedded in the biomaterial to advance cell therapies for myocardial infarction. The morphological details at the micrometre length scale resulted in a considerably higher and more heterogeneous mesh density.

The reconstruction of the biventricular FE model used a validated method developed in one of our previous studies (Sack *et al*. 2018). That study included an assessment of the mesh sensitivity, and although the mesh convergence analysis was not reported, mesh sensitivity guidelines for biventricular models were established. For quadratic tetrahedral elements, a mesh density of approximately 40,000 elements was sufficient to reach convergence of stress, strain and hemodynamic performance. The model used in the current study exceeds 200,000 elements, a deliberately fine resolution mesh, and a mesh sensitivity assessment was not deemed necessary.

Different strain energy density functions were used for the discrete representation of myocardial and injectate regions, contrary to studies that used the homogenization approach that combines the myocardium and biomaterial injectate (Wall *et al*. 2006, Wenk *et al*. 2011). This approach allowed capturing the distinctly different mechanical properties of the myocardium and injected biomaterial.

The current study adopted passive myocardial material parameters from Dương *et al*. (2018), who developed and validated the constitutive model for infarcted rat myocardium with experimental data from biaxial tension and uniaxial compression (Sirry *et al*. 2016). The active contraction in the healthy myocardial regions was modelled with a validated time-varying elastance model (Guccione *et al*. 1993). The parameters estimated by Guccione *et al*. (1993) have been used in several computational studies involving rat (Wise *et al*. 2016) and porcine models (Sack *et al*. 2018).

Therapeutic intramyocardial biomaterial injectates with different mechanical properties have been reported to induce different mechanical and functional responses in the infarcted heart (Wall *et al*. 2006).

The current study showed that an increasing elastic modulus (i.e. stiffness) of the biomaterial injectate reduces the ES and ED myocardial strains in the myofibre and cross-fibre directions in the healthy and infarcted regions of the BV geometry. This finding suggests that an increasing elastic modulus of the injectate for MI treatment may result in a beneficial effect of reducing the myocardial strains in the remote healthy regions of the BV geometry.

The biomaterial injectate is often used as a scaffold for therapeutically delivered cells for MI treatment. Thus, investigating the mechanical response of the biomaterial injectate delivered to the infarct is crucial for embedded cells’ mechanics and associated signalling. The current study showed that the mechanical response of the injectate depends on its stiffness. The maximum and minimum principal strains decreased for an increasing injectate elastic modulus. The upper range of the injectate’s elastic modulus (i.e. E_inj_ > 738 kPa) used in the study is unrealistic for PEG hydrogels and other injectable biomaterials (Rizzi *et al*. 2006, Singelyn *et al*. 2012). However, the current investigations aimed to explore the injectate response for an extensive range of injectate stiffness, and it was shown that the injectate deformations were negligible for these unrealistically high values of the injectate elastic modulus.

Limitations of the current study include the simplifying assumption of the *ex vivo* cardiac geometry reconstructed from μCT data as unloaded reference configuration for strain quantification.

The continuous mesh used for the myocardium and biomaterial injectate resulted in tied contact properties at the domain interfaces. More realistic contact interactions between myocardium and biomaterial will improve the simulation of the mechanics of the interface and the injectate but require experimental data that currently do not exist.

Considering the impact of the injectate stiffness on the ED and ES time point may improve the model’s accuracy. Furthermore, additional parameters such as injectate location and different injectate patterns may be considered to investigate their effect on the infarcted heart.

A linearly increasing pressure load on the cavity surfaces was used to implement the passive filling. This simple approach can be improved by using a more comprehensive representation of the circulatory system, including systemic arteries and veins, a pulmonary circuit, and the heart’s upper chambers and valves, as described by Sack *et al*. (2018).

The model’s anatomical details can be enhanced by implementing fibre orientation in the subject-specific biventricular geometry based on diffusion tensor magnetic resonance imaging.

## 5. Conclusions

This is the first computational study that generated and used a high-resolution microstructurally geometry of an *in situ* biomaterial injectate delivered one week after infarct induction in a biventricular cardiac finite element model of a rat heart with antero-apical infarct in the left ventricular free wall. With the microstructurally detailed in situ injectate geometry, the computational model of an infarcted rat heart offers vast potential for *in silico* studies of mechanotransduction and therapeutic signalling of cells transplanted with deliverable biomaterials in the infarcted heart, an animal model that has been extensively used in preclinical research of myocardial infarction.

## Nomenclature

Symbol: Description
a: Material parameter, dimension of stress
a_fs_: Material parameter defining coupling from in the fibre and sheet directions, with a dimension of stress
a_i_: Material parameter, defined for i = f and s in the fibre and sheet directions, respectively, with stress dimension
ā, ā_i_, ā_fs_: Govern the isotropic response of the infarcted myocardium
B: Governs the shape of peak isometric tension-sarcomere length relation
B: Dimensionless material parameter in Holzapfel model
b_fs_: Material parameter defining coupling from in the fibre and sheet directions, dimensionless
b_i_: Material parameter, defined for i = f and s in the fibre and sheet directions, respectively, dimensionless
C_10_: Coefficient used in Abaqus to describe the material stiffness in a Neo-Hookean strain energy density function
Ca_0_: Peak intracellular calcium concentration
D: Parameter for elastic materials defining the compressibility of the material
E: Elastic modulus
ECa_50_: Length-dependent calcium sensitivity
H: Parameter to define the pathological degree of the tissue
I_4f_: Transversely isotropic invariant in the fibre direction
I_4s_: Transversely isotropic invariant in the sheet direction
I_8fs_: Orthotropic invariant from coupling in fibre and sheet direction
I_i_: Isotropic invariants in principal directions
J: Third deformation gradient invariant as measures of the volume change of compressible materials
P: Parameter scaling the isotropic response of the diseased tissue
T: Stress tensor
T_(a)_: Active stress tensor
T_(p)_: Passive stress tensor
u_c_: Unit vector in the circumferential direction
u_l_: Unit vector in the longitudinal direction
W: Strain energy density function
α_h_: Helix angle
α_t_: Transverse angle
ε_EL_: Element strain; mean value of strains at integration points in an element
ε_IP,i_: Strain at integration point i in an element with i = 1 to 4
K: Bulk modulus
N: Poisson’s ratio
Σ: Stress

## Funding

This work was supported by financially supported by the National Research Foundation of South Africa (IFR14011761118 to TF), the South African Medical Research Council (SIR328148 to TF), and the CSIR Centre for High Performance Computing (CHPC Flagship Project Grant IRMA9543 to TF), and the Dr. Leopold und Carmen Ellinger Stiftung (UCT Three-Way PhD Global Partnership Programme Grant DAD937134 to TF). The funders had no role in study design, data collection and analysis, decision to publish, or preparation of the manuscript. Any opinion, findings, conclusions, and recommendations expressed in this publication are those of the authors, and therefore, the funders do not accept any liability.

## Conflict of Interests

Conflicts of interest do not exist.

## Data availability

Data supporting the results presented in this article are available on the University of Cape Town’s institutional data repository (ZivaHub) under http://doi.org/10.25375/uct.19630203 as YD Motchon, KL Sack, MS Sirry, E Pauwels, D Van Loo, A De Muynck, L Van Hoorebeke, NH Davies, T Franz. Effect of biomaterial stiffness on cardiac mechanics in a biventricular infarcted rat heart model with microstructural representation of in situ intramyocardial injectate. Cape Town, ZivaHub, 2022, DOI: 10.25375/uct.19630203.

## Notes

http://doi.org/10.25375/uct.19630203

### Competing Interest Statement

The authors have declared no competing interest.

### Summary of Updates

None

## References

Cai L, Wang Y, Gao H, Li Y, Luo X. A mathematical model for active contraction in healthy and failing myocytes and left ventricles. PLoS One 2017, 12(4): e0174834.

Chen J, Song S-K, Liu W, Mclean M, Allen JS, Tan J, Wickline SA, Yu X. Remodeling of cardiac fiber structure after infarction in rats quantified with diffusion tensor mri. American Journal of Physiology-Heart and Circulatory Physiology 2003, 285(3): H946–H54.

Christman KL, Fok HH, Sievers RE, Fang Q, Lee RJ. Fibrin glue alone and skeletal myoblasts in a fibrin scaffold preserve cardiac function after myocardial infarction. Tissue Eng 2004, 10(3-4): 403–9.

Costa KD, Takayama Y, Mcculloch AD, Covell JW. Laminar fiber architecture and three-dimensional systolic mechanics in canine ventricular myocardium. Am J Physiol 1999, 276(2): H595–607.

Dieudonné J-M. The left ventricle as confocal prolate spheroids. bulletin of mathematical biophysics 1969, 31(3): 433–9.

Duong MTN, Ach T, Alkassar M, Dittrich S, Leyendecker S. Numerical simulation of cardiac muscles in a rat biventricular model. 6th European Conference on Computational Mechanics (ECCM 6), 7th European Conference on Computational Fluid Dynamics (ECFD 7). UK, 2018.

Fan C, Shi J, Zhuang Y, Zhang L, Huang L, Yang W, Chen B, Chen Y, Xiao Z, Shen H, Zhao Y, Dai J. Myocardial-infarction-responsive smart hydrogels targeting matrix metalloproteinase for on-demand growth factor delivery. Adv Mater 2019, 31(40): e1902900.

Finegold JA, Asaria P, Francis DP. Mortality from ischaemic heart disease by country, region, and age: Statistics from World Health Organisation and United Nations. Int J Cardiol 2013, 168(2): 934–45.

Guccione J, Mcculloch A. Mechanics of active contraction in cardiac muscle: Part i—constitutive relations for fiber stress that describe deactivation. Journal of Biomechanical Engineering 1993, 115(1): 72–81.

Guccione J, Waldman L, Mcculloch A. Mechanics of active contraction in cardiac muscle: Part ii— cylindrical models of the systolic left ventricle. Journal of Biomechanical Engineering 1993, 115(1): 82–90.

Holmes JW, Borg TK, Covell JW. Structure and mechanics of healing myocardial infarcts. Annu Rev Biomed Eng 2005, 7: 223–53.

Holmes JW, Nunez JA, Covell JW. Functional implications of myocardial scar structure. Am J Physiol 1997, 272(5 Pt 2): H2123–30.

Holzapfel GA, Ogden RW. Constitutive modelling of passive myocardium: A structurally based framework for material characterization. Philos Trans A Math Phys Eng Sci 2009, 367(1902): 3445–75.

Hood WB, Bianco JA, Kumar R, Whiting RB. Experimental myocardial infarction: Iv. Reduction of left ventricular compliance in the healing phase. The Journal of Clinical Investigation 1970, 49(7): 1316-23 %@ 0021-9738.

Johnson TD, Christman KL. Injectable hydrogel therapies and their delivery strategies for treating myocardial infarction. Expert opinion on drug delivery 2013, 10(1): 59–72.

Kadner K, Dobner S, Franz T, Bezuidenhout D, Sirry MS, Zilla P, Davies NH. The beneficial effects of deferred delivery on the efficiency of hydrogel therapy post myocardial infarction. Biomaterials 2012, 33(7): 2060–6.

Kaptoge S, Pennells L, De Bacquer D, Cooney MT, Kavousi M, Stevens G, Riley LM, Savin S, Khan T, Altay S, Amouyel P, Assmann G, Bell S, Ben-Shlomo Y, Berkman L, Beulens JW, Björkelund C, Blaha M, Blazer DG, Bolton T, Bonita Beaglehole R, Brenner H, Brunner EJ, Casiglia E, Chamnan P, Choi Y-H, Chowdry R, Coady S, Crespo CJ, Cushman M, Dagenais GR, D’agostino Sr RB, Daimon M, Davidson KW, Engström G, Ford I, Gallacher J, Gansevoort RT, Gaziano TA, Giampaoli S, Grandits G, Grimsgaard S, Grobbee DE, Gudnason V, Guo Q, Tolonen H, Humphries S, Iso H, Jukema JW, Kauhanen J, Kengne AP, Khalili D, Koenig W, Kromhout D, Krumholz H, Lam TH, Laughlin G, Marín Ibañez A, Meade TW, Moons KGM, Nietert PJ, Ninomiya T, Nordestgaard BG, O’donnell C, Palmieri L, Patel A, Perel P, Price JF, Providencia R, Ridker PM, Rodriguez B, Rosengren A, Roussel R, Sakurai M, Salomaa V, Sato S, Schöttker B, Shara N, Shaw JE, Shin H-C, Simons LA, Sofianopoulou E, Sundström J, Völzke H, Wallace RB, Wareham NJ, Willeit P, Wood D, Wood A, Zhao D, Woodward M, Danaei G, Roth G, Mendis S, Onuma O, Varghese C, Ezzati M, Graham I, Jackson R, Danesh J, Di Angelantonio E. World health organization cardiovascular disease risk charts: Revised models to estimate risk in 21 global regions. The Lancet Global Health 2019, 7(10): e1332–e45.

Kortsmit J, Davies NH, Miller R, Macadangdang JR, Zilla P, Franz T. The effect of hydrogel injection on cardiac function and myocardial mechanics in a computational post-infarction model. Comput Methods Biomech Biomed Engin 2013a, 16(11): 1185–95.

Kortsmit J, Davies NH, Miller R, Zilla P, Franz T. Computational predictions of improved of wall mechanics and function of the infarcted left ventricle at early and late remodelling stages: Comparison of layered and bulk hydrogel injectates. Advances in Biomechanics and Applications 2013b, 1(1): 41–55.

Masschaele BC, Cnudde V, Dierick M, Jacobs P, Van Hoorebeke L, Vlassenbroeck J. Ugct: New x-ray radiography and tomography facility. Nuclear Instruments and Methods in Physics Research Section A: Accelerators, Spectrometers, Detectors and Associated Equipment 2007, 580(1): 266–9.

Mendis S, Puska P, Norrving B. Global atlas on cardiovascular disease prevention and control, World Health Organization, 2011.

Mininni S, Diricatti G, Vono MC, Giglioli C, Margheri M, Olivo G, Gensini G, Galanti G. Noninvasive evaluation of right ventricle systolic pressure during dynamic exercise by saline-enhanced doppler echocardiography in progressive systemic sclerosis. Angiology 1996, 47(5): 467–74.

Opie LH, Mayosi BM. Cardiovascular disease in sub-saharan africa. Circulation 2005, 112(23): 3536–40.

Pacher P, Mabley JG, Liaudet L, Evgenov OV, Marton A, Hasko G, Kollai M, Szabo C. Left ventricular pressure-volume relationship in a rat model of advanced aging-associated heart failure. Am J Physiol Heart Circ Physiol 2004, 287(5): H2132–7.

Reichek N, Devereux RB. Reliable estimation of peak left ventricular systolic pressure by m-mode echographic-determined end-diastolic relative wall thickness: Identification of severe valvular aortic stenosis in adult patients. Am Heart J 1982, 103(2): 202–3.

Rizzi SC, Ehrbar M, Halstenberg S, Raeber GP, Schmoekel HG, Hagenmüller H, Müller R, Weber FE, Hubbell JA. Recombinant protein-co-peg networks as cell-adhesive and proteolytically degradable hydrogel matrixes. Part ii: Biofunctional characteristics. Biomacromolecules 2006, 7(11): 3019–29.

Sack KL, Aliotta E, Ennis DB, Choy JS, Kassab GS, Guccione JM, Franz T. Construction and validation of subject-specific biventricular finite-element models of healthy and failing swine hearts from high-resolution dt-mri. Front Physiol 2018, 9(539): 539.

Sepantafar M, Maheronnaghsh R, Mohammadi H, Rajabi-Zeleti S, Annabi N, Aghdami N, Baharvand H. Stem cells and injectable hydrogels: Synergistic therapeutics in myocardial repair. Biotechnol Adv 2016, 34(4): 362–79.

Silveira-Filho LM, Coyan GN, Adamo A, Luketich SK, Menallo G, D’amore A, Wagner WR. Can a biohybrid patch salvage ventricular function at a late time point in the post-infarction remodeling process? Basic to Translational Science 2021, 6(5): 447–63.

Singelyn JM, Sundaramurthy P, Johnson TD, Schup-Magoffin PJ, Hu DP, Faulk DM, Wang J, Mayle KM, Bartels K, Salvatore M, Kinsey AM, Demaria AN, Dib N, Christman KL. Catheter-deliverable hydrogel derived from decellularized ventricular extracellular matrix increases endogenous cardiomyocytes and preserves cardiac function post-myocardial infarction. J Am Coll Cardiol 2012, 59(8): 751–63.

Sirry MS. Computational biomechanics of acute myocardial infarction and its treatments. Human Biology, Univerity of Cape Town, 2015, PhD: 145.

Sirry MS, Butler JR, Patnaik SS, Brazile B, Bertucci R, Claude A, Mclaughlin R, Davies NH, Liao J, Franz T. Characterisation of the mechanical properties of infarcted myocardium in the rat under biaxial tension and uniaxial compression. J Mech Behav Biomed Mater 2016, 63: 252–64.

Takayama Y, Costa KD, Covell JW. Contribution of laminar myofiber architecture to load-dependent changes in mechanics of lv myocardium. Am J Physiol Heart Circ Physiol 2002, 282(4): H1510–20.

Tyberg JV, Forrester JS, Wyatt HL, Goldner SJ, Parmley WW, Swan HJ. An analysis of segmental ischemic dysfunction utilizing the pressure-length loop. Circulation 1974, 49(4): 748–54.

Villarreal FJ, Lew WY, Waldman LK, Covell JW. Transmural myocardial deformation in the ischemic canine left ventricle. Circ Res 1991, 68(2): 368–81.

Vlassenbroeck J, Dierick M, Masschaele B, Cnudde V, Van Hoorebeke L, Jacobs P. Software tools for quantification of x-ray microtomography at the ugct. Nuclear Instruments and Methods in Physics Research Section A: Accelerators, Spectrometers, Detectors and Associated Equipment 2007, 580(1): 442–5.

Waldman LK, Nosan D, Villarreal F, Covell JW. Relation between transmural deformation and local myofiber direction in canine left ventricle. Circulation research 1988, 63(3): 550–62.

Walker JC, Ratcliffe MB, Zhang P, Wallace AW, Fata B, Hsu EW, Saloner D, Guccione JM. Mri-based finite-element analysis of left ventricular aneurysm. Am J Physiol Heart Circ Physiol 2005, 289(2): H692–700.

Wall ST, Walker JC, Healy KE, Ratcliffe MB, Guccione JM. Theoretical impact of the injection of material into the myocardium: A finite element model simulation. Circulation 2006, 114(24): 2627–35.

Wang H, Rodell CB, Lee ME, Dusaj NN, Gorman JH, 3rd, Burdick JA, Gorman RC, Wenk JF. Computational sensitivity investigation of hydrogel injection characteristics for myocardial support. J Biomech 2017, 64: 231–5.

Wenk JF, Eslami P, Zhang Z, Xu C, Kuhl E, Gorman Iii JH, Robb JD, Ratcliffe MB, Gorman RC, Guccione JM. A novel method for quantifying the in-vivo mechanical effect of material injected into a myocardial infarction. The Annals of Thoracic Surgery 2011, 92(3): 935–41.

Wenk JF, Wall ST, Peterson RC, Helgerson SL, Sabbah HN, Burger M, Stander N, Ratcliffe MB, Guccione JM. A method for automatically optimizing medical devices for treating heart failure: Designing polymeric injection patterns. J Biomech Eng 2009, 131(12): 121011.

Wise P, Davies NH, Sirry MS, Kortsmit J, Dubuis L, Chai CK, Baaijens FP, Franz T. Excessive volume of hydrogel injectates may compromise the efficacy for the treatment of acute myocardial infarction. International journal for numerical methods in biomedical engineering 2016, 32(12): e02772.

