## Supplemental tables for "Effect of biomaterial stiffness on cardiac mechanics in a biventricular infarcted rat heart model with microstructural representation of *in situ* intramyocardial injectate"

Supplemental Table 1. Constitutive parameters for passive mechanical behaviour of myocardium (Sack *et al.* 2018)

| Parameter | Value | Description |
| --- | --- | --- |
| a | 0.2065 kPa | Governs the isotropic response |
| b | 7.61 | Governs the isotropic response |
| a_f_ | 0.68 kPa | Governs additional stiffness in the fibre direction |
| b_f_ | 14.61 | Governs additional stiffness in the fibre direction |
| a_s_ | 0.945 kPa | Governs additional stiffness in the sheet direction |
| b_s_ | 12.67 | Governs additional stiffness in the sheet direction |
| a_fs_ | 5.55 x 10^-2^ kPa | Governs coupling stiffness in the fibre and sheet direction |
| b_fs_ | 3.12 | Governs coupling stiffness in the fibre and sheet direction |
| D | 0.2 x 10^-3^ kPa^-1^ | Defines the material incompressibility. Inversely proportional to the bulk modulus (K=2/D, defining the material’s resistance to compression) |
| h | 0-1 | Governs the health of the myocardium material point |
| p | 4.56 | Adjusts the passive response according to the stage of the infarct |

Supplemental Table 2. Constitutive parameters for active contraction in the myocardium (Guccione *et al.* 1993, Sack *et al.* 2018)

| Parameter | Value & unit | Description |
| --- | --- | --- |
| T_max_ | 60 kPa | Constitutive law scaling factor |
| Ca_0_ | 4.35 µmol/l | Peak intercellular calcium concentration |
| Ca_0max_ | 4.35 µmol/l | Maximum intercellular calcium concentration |
| B | 4,750 mm^-1^ | Governs the shape of the peak isometric tension-sarcomere length relation |
| l_0_ | 1.58 x 10^-3^ mm | The sarcomere length below which no active force develops |
| t_0_ | 0.15 s | Time to reach peak tension |
| m | 1,048.9 s/mm | Govern the shape of the linear relaxation duration and sarcomere relaxation |
| b | 1.5 s | Govern the shape of the linear relaxation duration and sarcomere relaxation |
| E_ff_ | - | Lagrangian strain tensor component aligned with the local muscle fibre direction |
| l_r_ | 2.03 x 10^-3^ mm | Initial sarcomere length |
